# Stochastic delays suppress noise in a genetic circuit with negative feedback

**DOI:** 10.1101/786491

**Authors:** Madeline Smith, Abhyudai Singh

## Abstract

We consider a mechanistic stochastic model of an autoregulatory genetic circuit with time delays. More specifically, a protein is expressed in random bursts from its corresponding gene. The synthesized protein is initially inactive and becomes active after a time delay. Rather than considering a deterministic delay, a key aspect of this work is to incorporate stochastic time delays, where delay is an independent and identically distributed random variable. The active protein inhibits its own production creating a negative feedback loop. Our analysis reveals that for an exponentially-distributed time delay, the noise in the protein levels decreases to the Poisson limit with increasing mean time delay. Interesting, for a gamma-distributed time delay contrasting noise behaviors emerge based on the negative feedback strength. At low feedback strengths the protein noise levels monotonically decreases to the Poisson limit with increasing average delay. At intermediate feedback strengths, the noise levels first increase to reach a maximum, and then decease back to the Poisson limit with increasing average delay. Finally, for strong feedbacks the protein noise levels monotonically increase with the average delay. For each of these scenarios we provide approximate analytical formulas for the protein mean and noises levels, and validate these results by performing exact Monte Carlo simulations. In conclusion, our results uncover a counter intuitive feature where inclusion of stochastic delays in a negative feedback circuit can play a beneficial role in buffering deleterious fluctuations in the level of a protein.

## I. INTRODUCTION

Gene-expression is the process by which a cell converts stored genetic information into protein molecules. These proteins are responsible for a variety of important cell functions. Like all biological systems, this process is based on a complex set of diffusion-driven biochemical reactions that are inherently random. Chemical systems generally operate with large numbers of molecules that average out the randomness. On the contrary, the key players in gene expression exist at very low copy numbers, amplifying the stochasticity inherent in biochemical processes [1]–[6]. For example, many mRNA species in *E. coli* and yeast are present at an average of 1 molecule or less per cell [2], [7], [8]. This stochasticity manifests as intercellular variation in the level of protein within an otherwise homogeneous cell population.

The prevalence of stochastic variation in gene product levels suggests that cells must exploit diverse regulatory mechanisms to either suppress it [9]–[14], or utilize it to their advantage via bet-hedging type strategies [15]–[19]. Perhaps the simplest example of buffering stochasticity is a negative feedback loop, where the protein directly or indirectly inhibits its own synthesis [20]–[35]. Such naturally-occurring feedbacks have been shown to be key motifs in gene regulatory networks [36]. Furthermore, design of in-vitro/in-silico feedback system for controlling gene expression are an intense area of current research [37]–[44].

A key question of interest is how feedback performance (in terms of noise control) is affected by time delays. While previous wok on this topic has assumed deterministic time delays [45], [46], we address this by considering stochastic delays in the feedback loop. These delays are mechanistically implemented through irreversible conversion reactions converting an inactive protein *X*_0_ into its active form *X_n_* via intermediates *X*_1_, … *X*_*n*–1_, and we refer to it as a *n*-step time delay. Assuming each conversion step occurs with a constant rate, it corresponds to a gamma-distributed time delay. For simplicity we will only consider *n* = 1 and 2 in this work. The negative feedback is implemented by having the active protein form *X_n_* suppress the production of *X*_0_.

Using both analytical approaches, such as the Linear Noise Approximation [47], [48], and exact Monte Carlo simulation we systematically study noise in the level of the active protein as a function of the average time-delay and the negative feedback strength. Our analysis reveals cases where inclusion of stochastic delays can reduce the protein noise level, and we identify parameter regimes associated with delay-mediated noise attenuation or enhancement. Finally, we quantify limits of noise suppression using negative feedback for a one-step and a two-step time delay.

The paper is organized as follows. In section II, we introduce a simple gene-expression model and obtain approximate analytical formulas for the protein mean and noise levels with and without negative feedback. In section III, we describe a one-step time delay model and show that increasing average time delay always suppresses noise to the Poisson limit. In section IV, we consider a two-step time delay model and find rich noise behaviours emerging for different feedback strengths. Finally, conclusions are presented in section V.

## II. MODEL WITHOUT TIME DELAY

### A. No Time Delay Model: Without Negative Feedback

We begin by considering a simple gene-expression model depicted in Fig. 1A. As shown experimentally across cell types [50]–[60], we consider a gene that synthesizes protein molecules in bursts. The burst events arrive as per a Poisson process with rate *k_x_*, and each event creates *B* number of protein molecules, where *B* is a discrete random variable drawn from the probability distribution

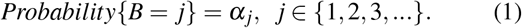

**Fig. 1:**
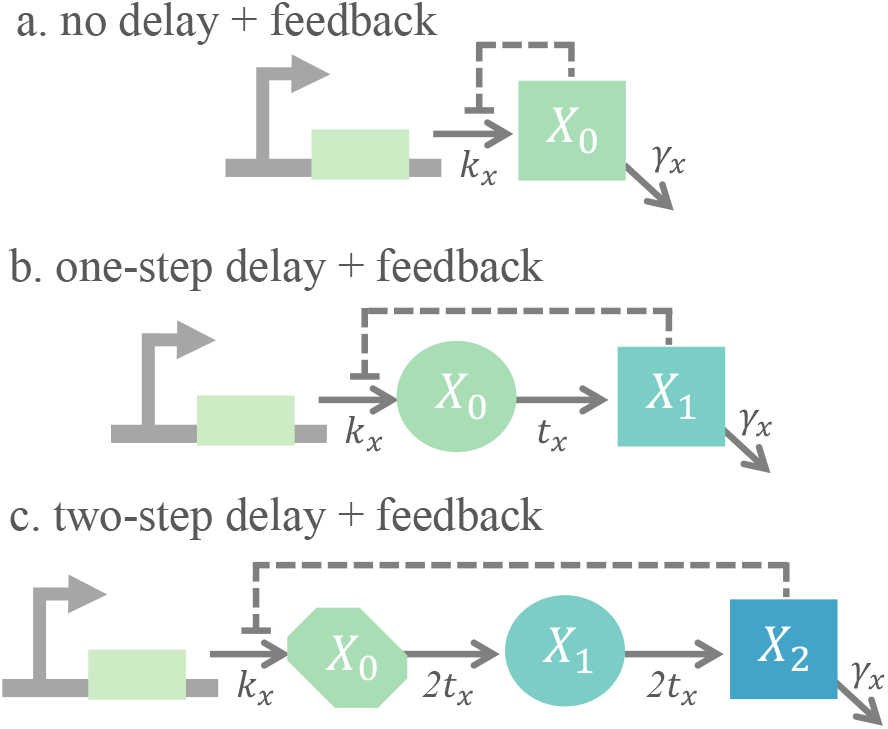
The gene expression process with feedback regulation and delay. A. Model of gene-expression with negative feedback that does not include a time delay. The gene promoter produces a protein *X*_0_ that inhibits its own production. B. Model of a one-step delay with negative feedback. The gene produces a protein *X*_0_ that undergoes a conversion reaction into an active protein species *X*_1_. The downstream protein, depicted as a square, inhibits the production of *X*_0_. C. Model of a two-step delay with negative feedback. The downstream protein, *X*_2_ square shaped, becomes active after two conversion reactions. The active protein inhibits he production of *X*_0_. In all three models, the active form of the protein degrades and the dashed line denotes negative feedback.

Moreover, each protein molecule degrades with a constant rate *γ_x_*. The protein species is denoted by *X*_0_ and its corresponding copy number at time *t* is denoted as ***x***_0_(*t*). The overall stochastic system is depicted in Table I and comprises of two events: protein production in bursts and protein degradation. When an event occurs, a corresponding reset in the population count ***x***_0_(*t*) is made, as listed in the middle column of Table I. Each event has a probability of occurrence in the next infinitesimal time interval (*t*,*t* + *dt*] that is listed in the rightmost column of Table I.

**TABLE I:**
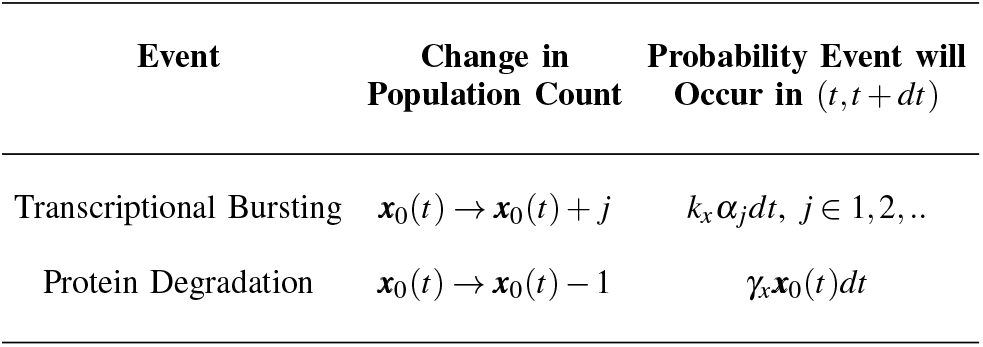
Stochastic model of gene expression with no feedbak and no delay.

| Event | Change in Population Count | Probability Event will Occur in $(t, t + dt)$ |
| --- | --- | --- |
| Transcriptional Bursting | $\mathbf{x}_0(t) \rightarrow \mathbf{x}_0(t) + j$ | $k_x \alpha_j dt, j \in 1, 2, \dots$ |
| Protein Degradation | $\mathbf{x}_0(t) \rightarrow \mathbf{x}_0(t) - 1$ | $\gamma_x \mathbf{x}_0(t) dt$ |

To derive an analytical expression for the noise in the protein level, we begin by writing differential equations describing the time evolution of the statistical moments of ***x***_0_(*t*). The time derivative of the expected value of any differentiable function *φ*(***x***_0_) is given by:

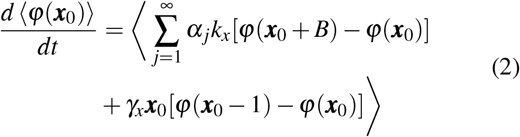

[61], [62], Here, and for the remainder of the paper, 〈.〉 is used to represent the expected value and 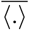 is used to represent the expected value at steady-state. By appropriately choosing *φ* we obtain the following moment dynamics:

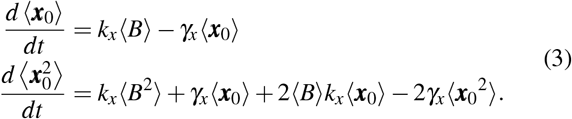

We quantify protein noise levels using the Fano factor, defined as the ratio of the variance to the mean. For a Poisson process, the variance is equal to the mean, thus the Fano factor is equal to one. Solving for the steady-state moments from (3) we obtain the following Fano factor

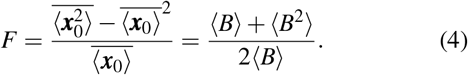

for a bursty gene expression model in the absence of any feedback regulation and delay. As expected, for non-bursty production *B* = 1 with a probability of 1, the Fano factor is equal to the Poisson limit.

### B. No Time Delay Model: Incorporating Negative Feedback

Next, negative feedback is implemented into the model by considering that the protein inhibits its own burst arrival rate. In the stochastic formulation, the original production rate *k_x_* becomes *k_x_*(***x***_0_), a monotonically decreasing function of the protein count ***x***_0_. Feedback implementation introduces nonlinearity in the model making the stochastic system analytically intractable. Assuming small fluctuations in the protein count around its set point, we linearize *k_x_*(***x***_0_) about the steady-state average number of protein molecules 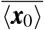

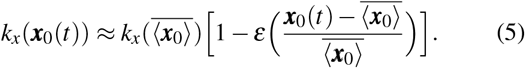

The dimensionless constant:

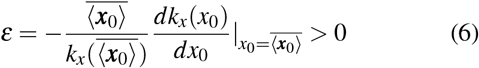

is the log sensitivity of the production rate to changes in ***x***_0_ and can be interpreted as the feedback strength [63].

As done in the previous section, we find the time evolution of the statistical moments. However, to implement the feedback, the approximated production rate *k_x_*(***x***_0_(*t*)) in (5) replaces *k_x_* in (2). By choosing *φ* as an appropriate monomial, we obtain the following moment dynamics:

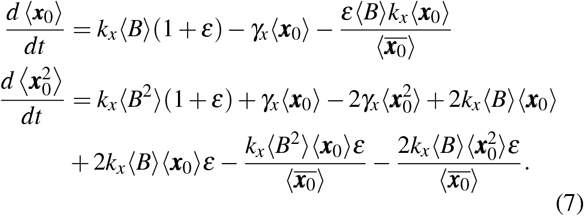

Solving for the steady-state moments yields the following Fano factor

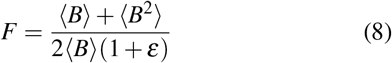

with negative feedback regulation, and increasing the feedback strength *ε* results in decreased noise.

## III. ONE-STEP TIME DELAY

### A. One-Step Time Delay Model: Without Negative Feedback

Next, we consider the effects of a stochastic time delay on the protein level noise. We begin by including a one-step time delay where the species *X*_0_ no longer degrades but instead undergoes a conversion reaction into the active protein form *X*_1_. This conversion occurs at a rate denoted by *t_x_*, and *X*_1_ degrades with a constant rate *γ_x_*. This one-step conversion corresponds to an exponentially-distributed delay with mean 1/*t_x_*, and we are interested in how this effects the noise levels of the final downstream active protein *X*_1_.

The stochastic formulation of the one-step delay model is shown in Table II, and comprises of three events: protein production, conversion, and degradation. Now, both species count, ***x***_0_(*t*) and ***x***_1_ (*t*) must be updated when an event occurs, as described by the reset map in Table II.

**TABLE II:** Stochastic model of gene expression with one-step delay and no feedback regulation.

| Event | Change in Population Count | Probability Event will Occur in $(t, t+dt)$ |
| --- | --- | --- |
| Transcriptional Bursting | $\mathbf{x}_0(t) \rightarrow \mathbf{x}_0(t) + j$ | $k_x \alpha_j dt, j \in 1, 2, \dots$ |
| One-Step Conversion | $\mathbf{x}_0(t) \rightarrow \mathbf{x}_0(t) - 1$<br>$\mathbf{x}_1(t) \rightarrow \mathbf{x}_1(t) + 1$ | $t_x \mathbf{x}_0(t) dt$ |
| Protein Degradation | $\mathbf{x}_1(t) \rightarrow \mathbf{x}_1(t) - 1$ | $\gamma_x \mathbf{x}_1(t) dt$ |

For the one-step delay model, the time-derivative of the expected value of any differentiable function *φ*(***x***_0_,***x***_1_) is given by:

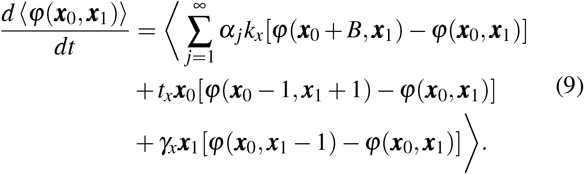

For appropriate choices for *φ*(***x***_0_,***x***_1_) we obtain the following moment dynamics:

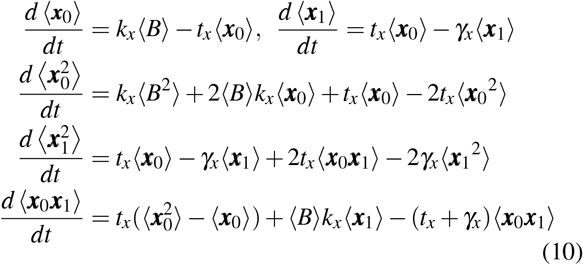

that result in the steady-state Fano factor

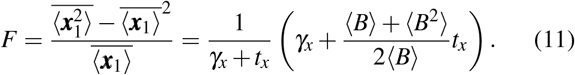

This Fano factor with a one-step delay is always smaller than the Fano factor without delay in (4), and with increasing mean delay

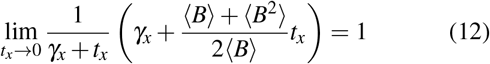

noise levels are reduced to the Poisson limit. Thus, stochastic delays function to effectively filter out underlying bursty production [64], [65].

### B. One-Step Time Delay: Incorporating Negative Feedback

Next, negative feedback is included into the one-step delay model by having the active protein *X*_1_ inhibit production of *X*_0_ via the burst arrival rate (Fig. 1B). In a similar fashion as in section II, we incorporate feedback by assuming that the production rate is a monotonically decreasing function *k_x_*(***x***_1_) of the active protein count *x*_1_. As before, *k_x_*(***x***_1_) is linearized about the steady-state protein level 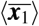

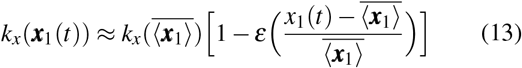

with the feedback strength given by

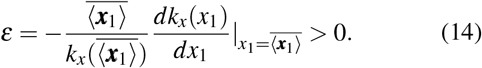

Next, the approximated production rate *k_x_*(***x***_1_(*t*)) in (13) is substituted in place of *k_x_* in (9). Repeating the procedure we again obtain the moment dynamics using (9) and solve it in the same manner as the previous section to find the Fano factor:

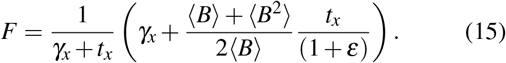

This formula provides two key insights. First, the Fano factor approaches a value of one with increasing mean delay (*t_x_* → 0) irrespective of the feedback strength *ε*. We illustrate this behavior in Figures 2, 3 and 4 where we further confirm these analytical predictions by performing exact Monte Carlo simulations using the Stochastic Simulation Algorithm [49]. Second, while the noise levels decrease with increasing feedback strength *ε* they approach a non-zero limit

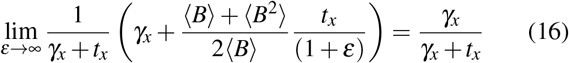

which represents a fundamental limit of noise suppression with a one-step time delay.

**Fig. 2:**
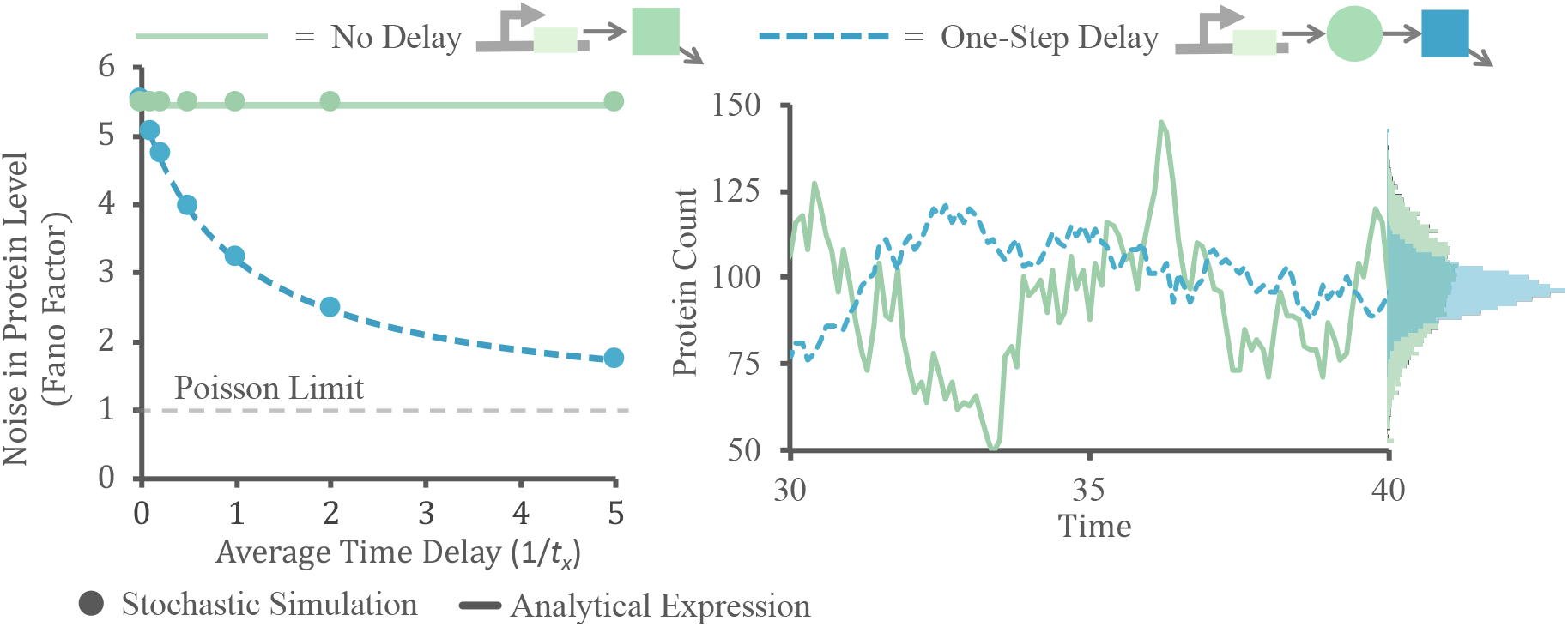
*Left:* Protein noise level plotted based on increasing mean time delay. The green-solid line denotes noise of the downstream protein in the no delay model. The blue-dashed line denotes noise of the downstream protein in the one-step delay model. As the delay is increased, and thus *t_x_* is decreased, the noise in the one-step delay becomes equal to 1. Solid circles represent noise values obtained by running a large number of Monte Carlo simulations. *Right:* Realizations of protein counts from stochastic simulation for the no delay (green solid), and the one-step delay model (blue dashed) using the Stochastic Simulation Algorithm [49]. The histogram for the protein level clearly reveals greater variation in the no delay model (green) compared to the one-step delay model (blue). The remaining parameters have the values: *B* = 10 with probability one, *k_x_* = 10, and *γ_x_* 1.

**Fig. 3:**
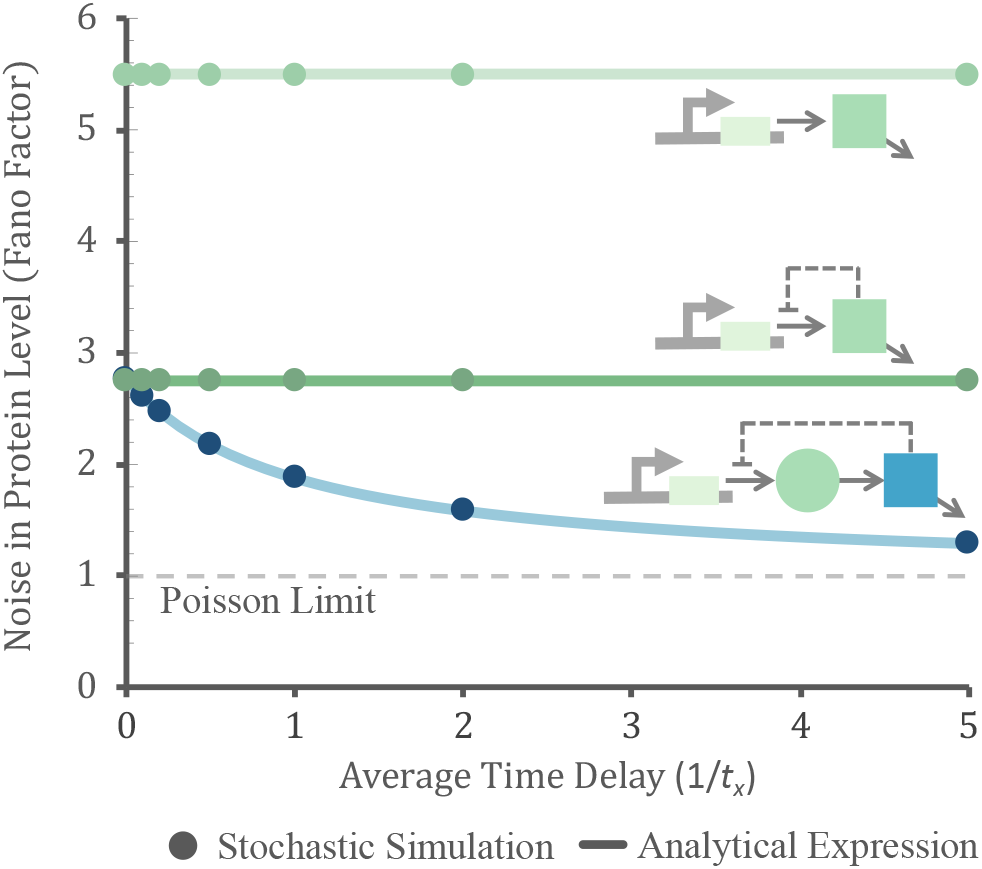
Noise in protein level is plotted as a function of increasing average time delay to compare the no delay without feedback model (light green), no delay with feedback model (dark green), and the one-step delay with feedback model (blue). The one-step time delay model with feedback provides the greatest noise attenuation. As the average time delay is increased, the noise is reduced to the Poisson limit. The remaining parameters have the values: *ε* = 1, *B* = 10 with probability one, *k_x_* = 10, and *γ_x_* 1. Solid lines are prediction from analytical formulas, while circles are values obtained from Monte Carlo simulations.

**Fig. 4:**
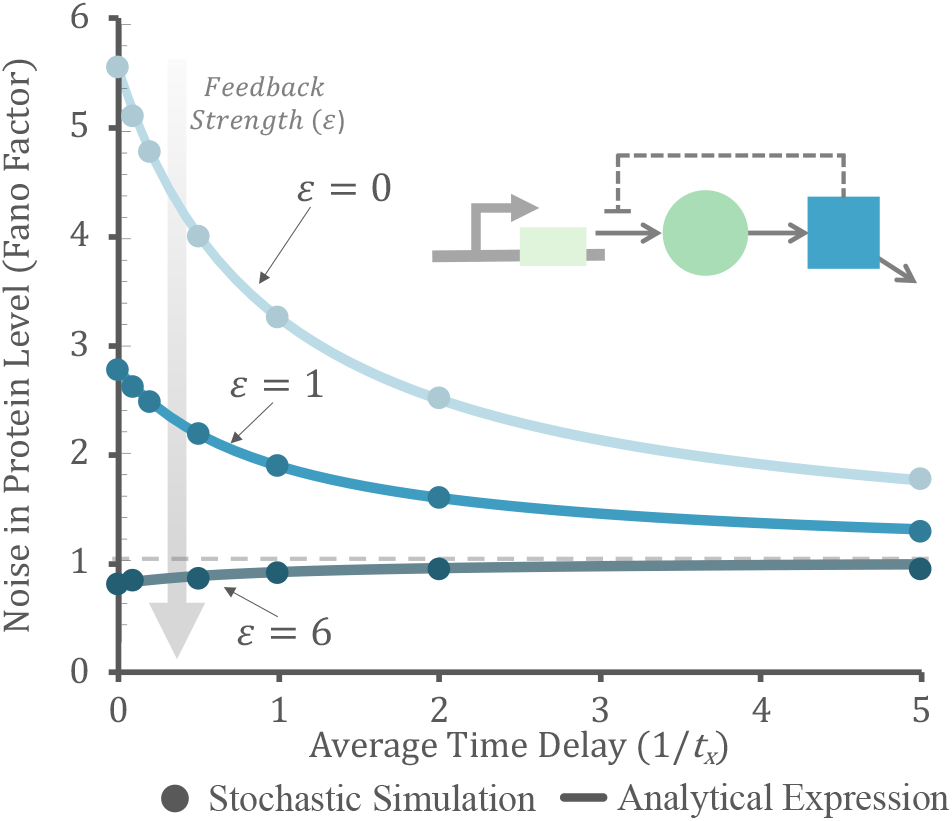
Protein noise level in a one-step delay model with different feedback strength. For all feedback strengths, as the time delay is increased, the noise converges to the Poisson limit. Note that if the initial noise is sub-Poissonian (Fano factor less than one), then noise increases with the average time delay. The remaining parameters have the values: *B* = 10 with probability one, *k_x_* = 10, and *γ_x_* = 1. Solid lines are prediction from analytical formulas, while circles are values obtained from Monte Carlo simulations.

## IV. Two-Step Time Delay

### A. Two-Step Time Delay Model: Without Negative Feedback

We further analyze stochastic time delays by considering a gene-expression circuit featuring a two-step delay. This circuit is illustrated in Fig. 1C. The model features three protein states: *X*_0_,*X*_1_,*X*_2_. In this set up, the gene produces *X*_0_, which is converted into *X*_1_ that is then converted into the active protein, *X*_2_. Each of the conversion reactions occur at a rate 2*t_x_* to maintain the average delay to be 1/*t_x_*. Only the active protein *X*_2_ degrades at a constant rate *γ_x_*. The overall stochastic model is shown in Table III, and using the corresponding extended generator

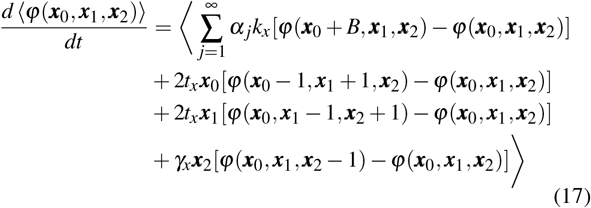

with appropriate choices for *φ*(***x***_0_,***x***_1_,***x***_2_) we obtain the following moment dynamics:

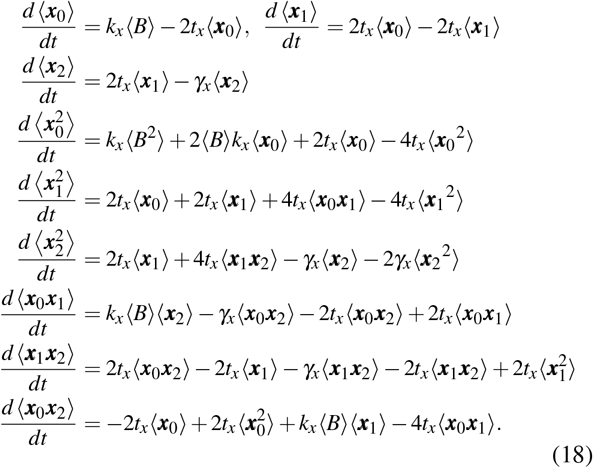

**TABLE III:**
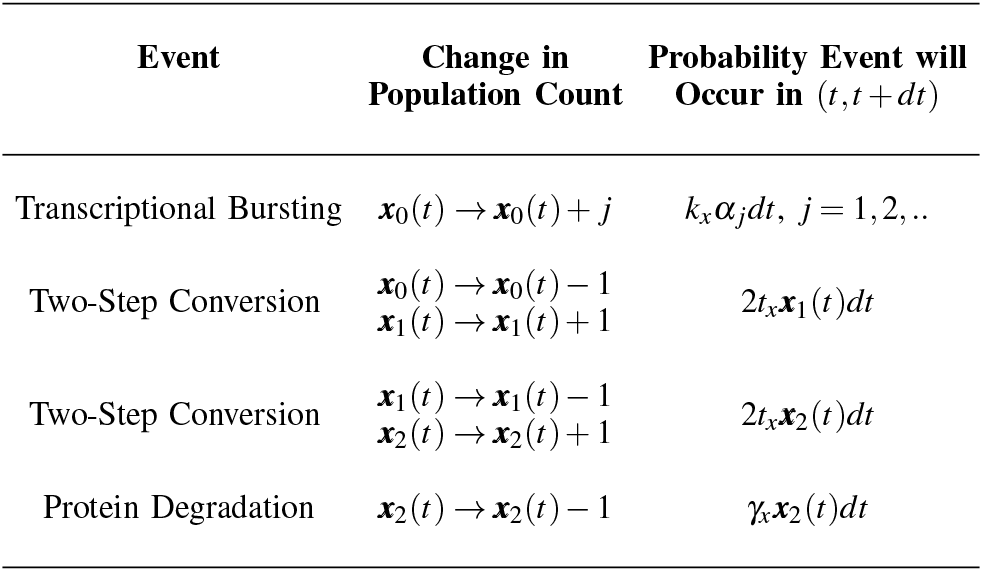
Stochastic gene expression model with a two-step time delay and no feedback regulation.

| Event | Change in Population Count | Probability Event will Occur in $(t, t + dt)$ |
| --- | --- | --- |
| Transcriptional Bursting | $\mathbf{x}_0(t) \rightarrow \mathbf{x}_0(t) + j$ | $k_x \alpha_j dt, j = 1, 2, \dots$ |
| Two-Step Conversion | $\mathbf{x}_0(t) \rightarrow \mathbf{x}_0(t) - 1$<br>$\mathbf{x}_1(t) \rightarrow \mathbf{x}_1(t) + 1$ | $2t_x \mathbf{x}_1(t) dt$ |
| Two-Step Conversion | $\mathbf{x}_1(t) \rightarrow \mathbf{x}_1(t) - 1$<br>$\mathbf{x}_2(t) \rightarrow \mathbf{x}_2(t) + 1$ | $2t_x \mathbf{x}_2(t) dt$ |
| Protein Degradation | $\mathbf{x}_2(t) \rightarrow \mathbf{x}_2(t) - 1$ | $\gamma_x \mathbf{x}_2(t) dt$ |

Solving for the steady-state moments, yields the Fano factor

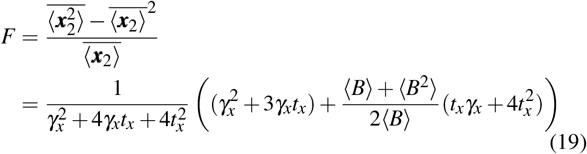

describing noise in the two-step delay without feedback. As expected, in the limit of small delay (*t_x_* → ∞) we recover the Fano factor without feedback (4), and the Fano factor reduces to the Poisson limit in the limit of large delay (*t_x_* → 0). It is important to point out that this noise level is higher than what we derived for a one-step delay in (11), but is smaller than the Fano factor without delay.

### B. Two-Step Time Delay: Incorporating Negative Feedback

Next, negative feedback is incorporated in the two-step time delay model by having *X*_2_ inhibit the production rate of *X*_0_. As done above, we approximate this production rate as:

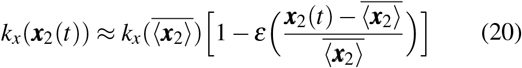

with feedback strength

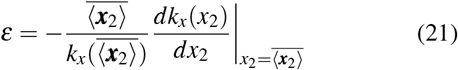

representing the log sensitivity of production rate *k_x_* (***x***_2_) to changes in ***x***_2_(*t*).

As done previously, replacing this approximated production rate *k_x_*(***x***_2_(*t*)) with *k_x_* in (17), and analyzing the subsequent moment dynamics yields

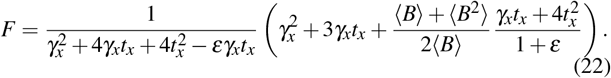

Before we interpret this formula, it is important to point out that with three species interlocked in a feedback loop, the corresponding deterministic dynamics can become unstable. A standard stability analysis of the mean dynamics reveals that the system is unstable for feedback strengths larger than

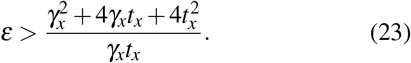

Since the right-hand-side of (23) is lower bounded by 8 (attains a minimum value when *t_x_* = *γ_x_*/2), *ε* > 8 is a necessary condition for unstable deterministic dynamics. Returning to the Fano factor we can see that as *ε* increases and approaches the stability boundary, the Fano factor becomes unbounded. In Fig. 5 we plot the Fano factor as a function of the mean time delay for different feedback strengths. Interestingly, contrasting behaviors emerge with noise monotonically decreasing to one (for low feedback strength), noise first increases to reach a maxima and then decreases to one (medium feedback strength), and noise monotonically increases to become unbounded (high feedback strength). Note in the later case there is a range of average time delays where the system is unstable, and as the average time delay is further increased the system again becomes stable and the Fano factor decreases to one.

**Fig. 5:**
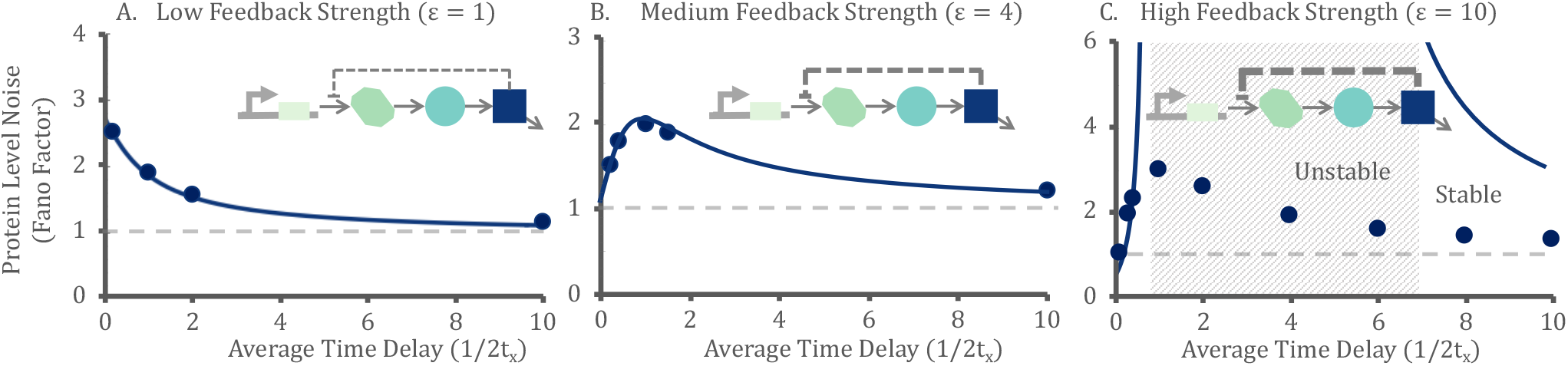
Protein noise levels with increasing average time delay for different feedback strengths. *A:* When feedback strength is low, noise monotonically decreases to 1. *B:* In the case of medium feedback strength, noise increases and then decreases to 1. *C*: When feedback strength is high, noise levels increases and become unbounded as the system becomes unstable. Based on stochastic simulations, noise follows the same qualitative behaviour as in the medium feedback scenario where it increases before decreasing to 1. The remaining parameters have the values: *B* = 10 with probability one, *k_x_* = 10, and *γ_x_* = 1. Solid lines are prediction from analytical formulas, while circles are values obtained from Monte Carlo simulations.

We next investigate how Fano factor varies with feedback strength for a fixed average time delay. Our results show that inclusion of feedback decreases noise levels (i.e. the sign of *dF*/*dε* is negative for low feedback strengths), but then Fano factor increases and becomes unbounded as the feedback becomes destabilizing. A detailed analysis of (22) reveals the lowest noise for a given two-step time delay to be

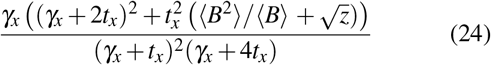

where

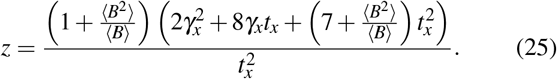

For example, if *B* = 1 with probability one, this noise suppression limit reduces to

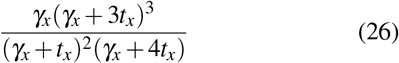

which monotonically decreases with increasing *t_x_* (i.e., smaller average delays).

## V. Conclusion

We have investigated the magnitude of stochastic fluctuations in protein counts subject to negative feedback regulation *with* and *without* stochastic delays in the feedback loop. When proteins are synthesized in bursts, the noise level is super-Poissonian (i.e., Fano factor larger than one) in the absence of both feedback and delay. Our results show that for a one-step delay, increasing the average time delay reduces the noise level to the Poissonian limit with and without negative feedback. If the protein production is non-bursty and feedback strength is high, then protein noise level may be sub-Poissonian initially, and in this case noise will increase to Poisson levels with increasing mean time delay. It is important to point out that for a one-step delay the corresponding deterministic system is always stable irrespective of the delay.

For a two-step delay the corresponding deterministic system can become unstable for sufficiently strong feedback (*ε* > 8). Our analysis reveals that for a two-step decay, noise reduces to the Poisson limit with increasing mean delay for weak feedback strengths. For strong yet stabilizing feedback (*ε* < 8), noise levels first increase, and then decrease back to the Poisson limit with increasing mean delay. Finally for strong and destabilizing feedback (*ε* > 8), noise levels increase with increasing mean delay and become unbounded as the system transitions from a stable to an unstable equilibrium. Note that while the Linear Noise Approximation predicts unbounded noise in the unstable regime, the actual noise levels as obtained by Monte Carlo simulations show a U-shape profile. Finally, as the mean delay is further increased, the deterministic system again becomes stable and noise levels attenuate to the Poisson limit.

### A. Future work

A natural area of future work is to investigate n-step stochastic delays for *n* ≥ 3 and explore the fundamental limit of noise suppression, and parameter regimes that lead to noise attenuation vs. amplification. It will also be interesting to investigate the stochastic dynamics in the regime where the corresponding deterministic system is unstable and exhibits a limit cycle. One approach here would be to linearize the nonlinear system around the deterministic limit cycle (and not the unstable equilibrium point), and then explore higher order statistical moments via moment dynamics. Moreover, one could compute auto-correlation functions to quantify the precision of oscillation as a function of protein bursting and stochastic delay.

## ACKNOWLEDGMENT

This work is supported by the National Science Foundation Grant ECCS-1711548.

## References

[1] W. J. Blake, M. Kaern, C. R. Cantor, and J. J. Collins, “Noise in eukaryotic gene expression,” Nature, vol. 422, pp. 633–637, 2003.

[2] A. Bar-Even, J. Paulsson, N. Maheshri, M. Carmi, E. O’ Shea, Y. Pilpel, and N. Barkai, “Noise in protein expression scales with natural protein abundance,” Nature Genetics, vol. 38, pp. 636–643, 2006.

[3] J. M. Raser and E. K. O’Shea, “Noise in gene expression: Origins, consequences, and control,” Science, vol. 309, pp. 2010–2013, 2005.

[4] A. Eldar and M. B. Elowitz, “Functional roles for noise in genetic circuits,” Nature, vol. 467, pp. 167–173, 2010.

[5] M. B. Elowitz, A. J. Levine, E. D. Siggia, and P. S. Swain, “Stochastic gene expression in a single cell,” Science, vol. 297, pp. 1183–1186, 2002.

[6] A. Raj and A. van Oudenaarden, “Nature, nurture, or chance: stochastic gene expression and its consequences,” Cell, vol. 135, pp. 216–226, 2008.

[7] Y. Taniguchi, P. J. Choi, G. W. Li, H. Chen, M. Babu, J. Hearn, A. Emili, and X. S. Xie, “Quantifying E. coli proteome and transcriptome with single-molecule sensitivity in single cells,” Science, vol. 329, pp. 533–538, 2010.

[8] J. R. S. Newman, S. Ghaemmaghami, J. Ihmels, D. K. Breslow, M. Noble, J. L. DeRisi, and J. S. Weissman, “Single-cell proteomic analysis of S. cerevisiae reveals the architecture of biological noise,” Nature Genetics, vol. 441, pp. 840–846, 2006.

[9] E. Libby, T. J. Perkins, and P. S. Swain, “Noisy information processing through transcriptional regulation,” Proceedings of the National Academy of Sciences, vol. 104, pp. 7151–7156, 2007.

[10] H. B. Fraser, A. E. Hirsh, G. Giaever, J. Kumm, and M. B. Eisen, “Noise minimization in eukaryotic gene expression,” PLOS Biology, vol. 2, p. e137, 2004.

[11] B. Lehner, “Selection to minimise noise in living systems and its implications for the evolution of gene expression,” Molecular Systems Biology, vol. 4, p. 170, 2008.

[12] R. Kemkemer, S. Schrank, W. Vogel, H. Gruler, and D. Kaufmann, “Increased noise as an effect of haploinsufficiency of the tumorsuppressor gene neurofibromatosis type 1 in vitro,” Proceedings of the National Academy of Sciences, vol. 99, pp. 13783–13788, 2002.

[13] D. L. Cook, A. N. Gerber, and S. J. Tapscott, “Modeling stochastic gene expression: implications for haploinsufficiency,” Proceedings of the National Academy of Sciences, vol. 95, pp. 15641–15646, 1998.

[14] A. Brock, H. Chang, and S. Huang, “Non-genetic heterogeneity – a mutation-independent driving force for the somatic evolution of tumours,” Nature Reviews Genetics, vol. 10, pp. 336–342, 2009.

[15] H. Maamar, A. Raj, and D. Dubnau, “Noise in gene expression determines cell fate in bacillus subtilis,” Science, vol. 317, pp. 526–529, 2007.

[16] S. M. Shaffer, M. C. Dunagin, S. R. Torborg, E. A. Torre, B. Emert, C. Krepler, M. Beqiri, K. Sproesser, P. A. Brafford, M. Xiao, E. Eggan, I. N. Anastopoulos, C. A. Vargas-Garcia, A. Singh, K. L. Nathanson, M. Herlyn, and A. Raj, “Rare cell variability and drug-induced reprogramming as a mode of cancer drug resistance,” Nature, vol. 546, pp. 431–435, 2017.

[17] N. Balaban, J. Merrin, R. Chait, L. Kowalik, and S. Leibler, “Bacterial persistence as a phenotypic switch,” Science, vol. 305, pp. 1622–1625, 2004.

[18] I. E. Meouche, Y. Siu, and M. J. Dunlop, “Stochastic expression of a multiple antibiotic resistance activator confers transient resistance in single cells,” Scientific Reports, vol. 6, p. 19538, 2016.

[19] R. Bahar, C. H. Hartmann, K. A. Rodriguez, A. D. Denny, R. A. Busuttil, M. E. Dolle, R. B. Calder, G. B. Chisholm, B. H. Pollock, C. A. Klein, and J. Vijg, “Increased cell-to-cell variation in gene expression in ageing mouse heart,” Nature, vol. 441, pp. 1011–1014, 2006.

[20] A. Becskei and L. Serrano, “Engineering stability in gene networks by autoregulation,” Nature, vol. 405, pp. 590–593, 2000.

[21] Y. Dublanche, K. Michalodimitrakis, N. Kummerer, M. Foglierini, and L. Serrano, “Noise in transcription negative feedback loops: simulation and experimental analysis,” Molecular Systems Biology, vol. 2, p. 41, 2006.

[22] A. Singh and J. P. Hespanha, “Optimal feedback strength for noise suppression in autoregulatory gene networks,” Biophysical Journal, vol. 96, pp. 4013–4023, 2009.

[23] D. Orrell and H. Bolouri, “Control of internal and external noise in genetic regulatory networks,” Journal of Theoretical Biology, vol. 230, pp. 301–312, 2004.

[24] A. Borri, P. Palumbo, and A. Singh, “The impact of negative feedback in metabolic noise propagation,” IET Systems Biology, pp. 179–186, 2016.

[25] I. Lestas, G. Vinnicombe, and J. Paulsson, “Fundamental limits on the suppression of molecular fluctuations,” Nature, vol. 467, pp. 174–178, 2010.

[26] Y. Tao, X. Zheng, and Y. Sun, “Effect of feedback regulation on stochastic gene expression,” Journal of Theoretical Biology, vol. 247, pp. 827–836, 2007.

[27] S. Kumar and A. J. Lopez, “Negative feedback regulation among sr splicing factors encoded by rbp1 and rbp1-like in drosophila,” The EMBO Journal, vol. 24, pp. 2646–2655, 2005.

[28] A. Singh and J. P. Hespanha, “Evolution of autoregulation in the presence of noise,” IET Systems Biology, vol. 3, pp. 368–378, 2009.

[29] D. J. Stekel and D. J. Jenkins, “Strong negative self regulation of prokaryotic transcription factors increases the intrinsic noise of protein expression,” BMC Systems Biology, 2008.

[30] H. El-Samad and M. Khammash, “Regulated degradation is a mechanism for suppressing stochastic fluctuations in gene regulatory networks,” Biophysical Journal, vol. 90, pp. 3749–3761, 2006.

[31] P. S. Swain, “Efficient attenuation of stochasticity in gene expression through post-transcriptional control,” Journal of Molecular Biology, vol. 344, pp. 956–976, 2004.

[32] M. Voliotis and C. G. Bowsher, “The magnitude and colour of noise in genetic negative feedback systems,” Nucleic Acids Research, 2012.

[33] R. Bundschuh, F. Hayot, and C. Jayaprakash, “The role of dimerization in noise reduction of simple genetic networks,” Journal of Theoretical Biology, vol. 220, pp. 261–269, 2003.

[34] J. M. Pedraza and J. Paulsson, “Effects of molecular memory and bursting on fluctuations in gene expression,” Science, vol. 319, pp. 339–343, 2008.

[35] P. Bokes, Y. Ting Lin, and A. Singh, “High cooperativity in negative feedback can amplify noisy gene expression,” Society for Mathematical Biology, vol. 80, pp. 1871–1899, 2018.

[36] U. Alon, “Network motifs: theory and experimental approaches,” Nature Reviews Genetics, vol. 8, pp. 450–461, 2007.

[37] R. Sawlekar, F. Montefusco, V. V. Kulkarni, and D. G. Bates, “Implementing nonlinear feedback controllers using dna strand displacement reactions,” IEEE Transactions, vol. 15, p. 443, 2016.

[38] A. Milias-Argeitis, S. Summers, J. Stewart-Ornstein, I. Zuleta, D. Pincus, H. El-Samad, M. Khammash, and J. Lygeros, “In silico feedback for in vivo regulation of a gene expression circuit,” Nature Biotechnology, vol. 29, 2011.

[39] E. Klavins, “Proportional-integral control of stochastic gene regulatory networks,” IEEE conference on Decision and Control, 2010.

[40] G. Buzi and M. Khammash, “Implementation considerations, not topological differences, are the main determinants of noise suppression properties in feedback and incoherent feedforward circuits,” PLoS Computational Biology, vol. 12, p. e1004958, 2016.

[41] J. Uhlendorf, A. Miermont, T. Delaveau, G. Charvin, F. Fages, S. Bottani, G. Batt, and P. Hersen, “Long-term model predictive control of gene expression at the population and single-cell levels,” IET Systems Biology, vol. 109, 2012.

[42] C. Briat, C. Zechner, and M. Khammash, “Design of a synthetic integral feedback circuit: Dynamic analysis and dna implementation,” ACS Synthetic Biology, vol. 5, 2016.

[43] C. Briat, A. Gupta, and M. Khammash, “Antithetic integral feedback ensures robust perfect adaptation in noisy biomolecular networks,” Cell Systems, vol. 2, 2016.

[44] M. E. Wall, W. S. Hlavacek, and M. A. Savageau, “Design principles for regulator gene expression in a repressible gene circuit,” Journal of Molecular Biology, vol. 332, pp. 861–876, 2003.

[45] E. Zavala and T. T. Marquez-Lago, “Delays induce novel stochastic effects in negative feedback gene circuits,” Biophysical Journal, pp. 467–478, 2014.

[46] M. Barrio, K. Burrage, A. Leier, and T. Tian, “Oscillatory regulation of hes1: Discrete stochastic delay modelling and simulation,” PLoS Comput Biol, vol. 2, p. e117, 2006.

[47] N. Van Kampen, Stochastic processes in physics and chemistry. Elsevier, 2011.

[48] S. Modi, M. Soltani, and A. Singh, “Linear noise approximation for a class of piecewise deterministic markov processes,” American Control Conference (ACC), 2018.

[49] D. T. Gillespie, “A general method for numerically simulating the stochastic time evolution of coupled chemical reactions,” Journal of Computational Physics, vol. 22, pp. 403–434, 1976.

[50] D. M. Suter, N. Molina, D. Gatfield, K. Schneider, U. Schibler, and F. Naef, “Mammalian genes are transcribed with widely different bursting kinetics,” Science, vol. 332, pp. 472–474, 2011.

[51] R. D. Dar, B. S. Razooky, A. Singh, T. V. Trimeloni, J. M. McCollum, C. D. Cox, M. L. Simpson, and L. S. Weinberger, “Transcriptional burst frequency and burst size are equally modulated across the human genome,” Proceedings of the National Academy of Sciences, vol. 109, pp. 17454–17459, 2012.

[62] A. Singh and J. P. Hespanha, “Approximate moment dynamics for chemically reacting systems,” IEEE Transactions on Automatic Control, vol. 56, pp. 414–418, 2011.

[52] T. Fukaya, B. Lim, and M. Levine, “Enhancer control of transcriptional bursting,” Cell, vol. 166, pp. 358–368, 2015.

[53] C. R. Bartman, S. C. Hsu, C. C.-S. Hsiung, A. Raj, and G. A. Blobel, “Enhancer regulation of transcriptional bursting parameters revealed by forced chromatin looping,” Molecular Cell, vol. 62, pp. 237–247, 2016.

[54] A. M. Corrigan, E. Tunnacliffe, D. Cannon, and J. R. Chubb, “A continuum model of transcriptional bursting,” eLife, vol. 5, p. e13051, 2016.

[55] S. Chong, C. Chen, H. Ge, and X. S. Xie, “Mechanism of transcriptional bursting in bacteria,” Cell, vol. 158, pp. 314–326, 2014.

[56] A. Singh, B. Razooky, C. D. Cox, M. L. Simpson, and L. S. Weinberger, “Transcriptional bursting from the HIV-1 promoter is a significant source of stochastic noise in HIV-1 gene expression,” Biophysical Journal, vol. 98, pp. L32–L34, 2010.

[57] R. D. Dar, S. M. Shaffer, A. Singh, B. S. Razooky, M. L. Simpson, A. Raj, and L. S. Weinberger, “Transcriptional bursting explains the noise–versus–mean relationship in mRNA and protein levels,” PLOS ONE, vol. 11, p. e0158298, 2016.

[58] I. Golding, J. Paulsson, S. Zawilski, and E. Cox, “Real-time kinetics of gene activity in individual bacteria,” Cell, vol. 123, pp. 1025–1036, 2005.

[59] A. Raj, C. S. Peskin, D. Tranchina, D. Vargas, and S. Tyagi, “Stochastic mRNA synthesis in mammalian cells,” PLOS Biology, vol. 4, p. e309, 2006.

[60] A. Singh, B. S. Razooky, R. D. Dar, and L. S. Weinberger, “Dynamics of protein noise can distinguish between alternate sources of geneexpression variability,” Molecular Systems Biology, vol. 8, p. 607, 2012.

[61] J. P. Hespanha and A. Singh, “Stochastic models for chemically reacting systems using polynomial stochastic hybrid systems,” International Journal of Robust and Nonlinear Control, vol. 15, pp. 669–689, 2005.

[63] A. Singh, “Negative feedback through mRNA provides the best control of gene-expression noise,” IEEE Transactions on Nanobioscience, vol. 10, pp. 194–200, 2011.

[64] A. Singh and P. Bokes, “Consequences of mRNA transport on stochastic variability in protein levels,” Biophysical Journal, vol. 103, pp. 1087–1096, 2012.

[65] T. Stoeger, N. Battich, and L. Pelkmans, “Passive noise filtering by cellular compartmentalization,” Cell, vol. 164, pp. 1151–1161, 2016.

